# Color Constant Representations in Early Visual Cortex

**DOI:** 10.1101/2022.06.01.494333

**Authors:** Anke Marit Albers, Elisabeth Baumgartner, Karl R. Gegenfurtner

## Abstract

The light entering our eyes is the product of the illumination and the surface reflectance of an object. Although this light changes considerably when the illumination changes, we are usually able to perceive objects as stable in color. To investigate how the brain achieves color constancy, we measured BOLD fMRI while 19 participants either observed patches of light that appear colored (yellow, blue) under a spectrally neutral illuminant, or spectrally neutral gray patches that appear colored under simulated blue and yellow illumination conditions. Under bluish illumination, the neutral gray patches appeared yellow; under yellowish illumination, the same gray patches appeared blue. We successfully trained a classifier to discriminate between the blue- and yellow-colored patches in V1-V4. Crucially, we then tested whether this same classifier could also distinguish between the apparent blue and yellow induced by the illuminants. The neural representations for apparent blue and yellow resembled colorimetric blue and yellow in V1, V3 and V4. A control experiment showed that apparent lightness cannot explain these effects. These findings suggest that not only colorimetric, but also apparent color is represented to some degree in retinotopic visual cortex, as early as in V1. Furthermore, a small frontal region, the Rolandic operculum, showed activation for apparent color, possibly playing a role in color constancy.

## Introduction

Color is an important aspect of processing in the human visual system, starting with three different types of cone photoreceptors in the retina and continuing throughout the visual pathways (see reviews by Conway, 2009, 2014; Conway et al., 2010; Gegenfurtner & Kiper, 2003). However, color is strictly speaking not a property of the objects that we perceive. While the reflecting properties of objects are stable and invariant, the light that is reflected from the object and that enters our eyes is also dependent on the illumination. Therefore, the spectral and chromatic properties change under varying illuminations. Yet, humans usually perceive the color of objects as being invariant under different illuminants (for reviews, see Foster, 2011; Hurlbert, 2007; Maloney, 1999; Smithson, 2005). This allows us to recognize objects in different circumstances, categorize colors consistently and communicate them reliably (Witzel and Gegenfurtner, 2018). As a side effect, the same colorimetric input can be perceived differently if it appears in scenes that seem consistent with different illuminations (for examples, see e.g. (Gegenfurtner, 2003; Land, 1977; Wachtler et al., 2003; Witzel and Gegenfurtner, 2018) and the work by Akiyoshi Kitaoka: http://www.psy.ritsumei.ac.jp/~akitaoka/colorconstancy7e.html). A well-known example is the cube by Lotto and Purves (2004), where a colorimetrically similar patch appears markedly different under a simulated blue or yellow illuminant. In this case colorimetry – the light entering the eye reflected in retinal cone absorptions – and (subjective) color appearance are dissociated from each other as an effect of color constancy calculations.

Color constancy is known to rely on a multitude of cues. Some of them are relatively easy to compute, such as the global color within a scene (“gray world assumption”), or the cone contrasts across edges in a scene (Conway et al., 2007, 2002; Conway and Livingstone, 2006; Foster et al., 2006; Kentridge et al., 2007, 2004; Shapley and Hawken, 2002). Others are much more complex, such as inter-reflections between different parts of a scene (Bloj et al., 1999). The human visual system seems to combine information from these different cues (Hansen et al., 2007; Kraft and Brainard, 1999; Weiss et al., 2017) to find a representation of the world that is invariant of illumination. The more realistic experimental scenarios are, the better the degree of invariance that is achieved (Foster, 2011; Witzel and Gegenfurtner, 2018).

Presumably, the computations underlying the above mechanisms are performed at different stages of information processing in the human eye and brain. While some adaptation can even take place in the cone photoreceptors, edge contrast is presumably first computed in V1, and for a more global normalization larger receptive fields are necessary, as they are for example found in V4. Currently, not much is known about the transition from colorimetry-based to appearance-based representations of color in the brain. Early work reported color constant responses in cells in V4 (Schein and Desimone, 1990; Zeki, 1983). Patient studies imply that lesions of the medial and lateral posterior fusiform gyrus, which corresponds to human V4 (Lueck et al., 1989), lead to color constancy impairments (Clarke et al., 1998; Kennard et al., 1995; Walsh, 1999; Zeki et al., 1999). A selective impairment for color constancy in humans has been linked with lesions in higher cortical regions (Rüttiger et al., 1999), but the output from these higher order regions could be fed back to influence representations in striate cortex.

More recent studies in primates suggest that cells as early as V1 can respond in a color constant fashion (Wachtler et al., 2003, 2001). However, whether human V1 also contains color-constant representations is not yet clear. A patient with a lesion in occipito-parietal cortex, but spared V1 and V2, was aware of colors, but did not show color constancy (Zeki et al., 1999). Another patient with a removed right striate cortex was unable to discriminate colors, yet showed some spared color constancy (Heywood et al., 1991), although he was unable to use chromatic contrasts (Kentridge et al., 2004), discounting V1 as the most important source of constancy. A recent fMRI study showed that for complex scenes, there were surface color representations across illuminations in V4α and V1 (Bannert and Bartels, 2017). Human V1 has also been implicated in the representation of memory colors (Bannert and Bartels, 2013) a process that also involves color appearance rather than colorimetric representations.

Here we asked to which degree the neural coding in human early visual cortex are color constant – that is, appearance based and invariant to illuminant changes – and to what extent it is determined by the photoreceptor activations in the eye. We used simplified versions of the cube by Lotto & Purves (2004), which allowed a dissociation between the neural representations of colorimetry and color appearance. Although the surfaces of the center squares in the stimuli were colorimetrically neutral, they induced a blue appearance under the yellow illuminant and yellow appearance under the blue illuminant.

Participants observed both these colorimetrically neutral stimuli, as well as stimuli with colorimetrically blue and yellow surfaces, while we measured brain activity using fMRI. We subsequently employed Multi-Voxel Pattern Analyses (MVPA) to test whether a classifier trained on colorimetric blue and yellow, could also discriminate between brain activity patterns for inferred blue and yellow, specifically in V1-V4, suggesting representations of surface color invariant to illuminant changes. In a control experiment, we used grayscale versions of the stimuli to test for the effect of lightness in the color induction effect. Our results suggest that color constant representations appear in several visual areas and as early as V1 in the visual hierarchy.

## Methods

### 2.1 Participants & Procedure

Nineteen participants (12 female, ages 19-28, mean age 23.5 years) with normal or corrected-to-normal vision were recruited from the student population at the Justus Liebig University Giessen. Participants were first screened for normal color vision using the Ishihara plates (Ishihara, 2004). They then participated in up to three fMRI sessions. All participants performed the first session with the main color induction experiment that contained the apparent color condition and the colorimetric color condition. All participants performed a second session with the functional localizers and retinotopy. For one participant retinotopy could not be drawn, leaving 18 participants for the decoding analyses. Most participants (N = 14) took part in a third session, in which they performed the grayscale control experiment. The exact order of these sessions varied somewhat due to practical considerations. The study was approved by the local ethics committee at the Psychology Department of Giessen University (approval number LEK 2017-0030), and all participants signed informed consent forms before participation. They received course credit or a monetary reward for their participation.

### 2.2 Colors and stimuli

#### 2.2.1 Apparent Color Condition

Our stimuli were inspired by the cube illusion introduced by Lotto and Purves (2004; http://purveslab.net/see-for-yourself). Their 3D cube consisted of colored patches rendered under simulated bluish and yellowish illuminants, where the center patch had the same neutral physical wavelength spectrum (a colorimetrically neutral gray), yet appeared to be of different color under the two different illuminants (Figure 1C): under the bluish illumination the center patch appeared yellow; under the yellowish illuminant it appeared blue. We derived the RGB values of the surrounding patches and the illuminated background from the original demo by Lotto & Purves (2004).

**Figure 1:**
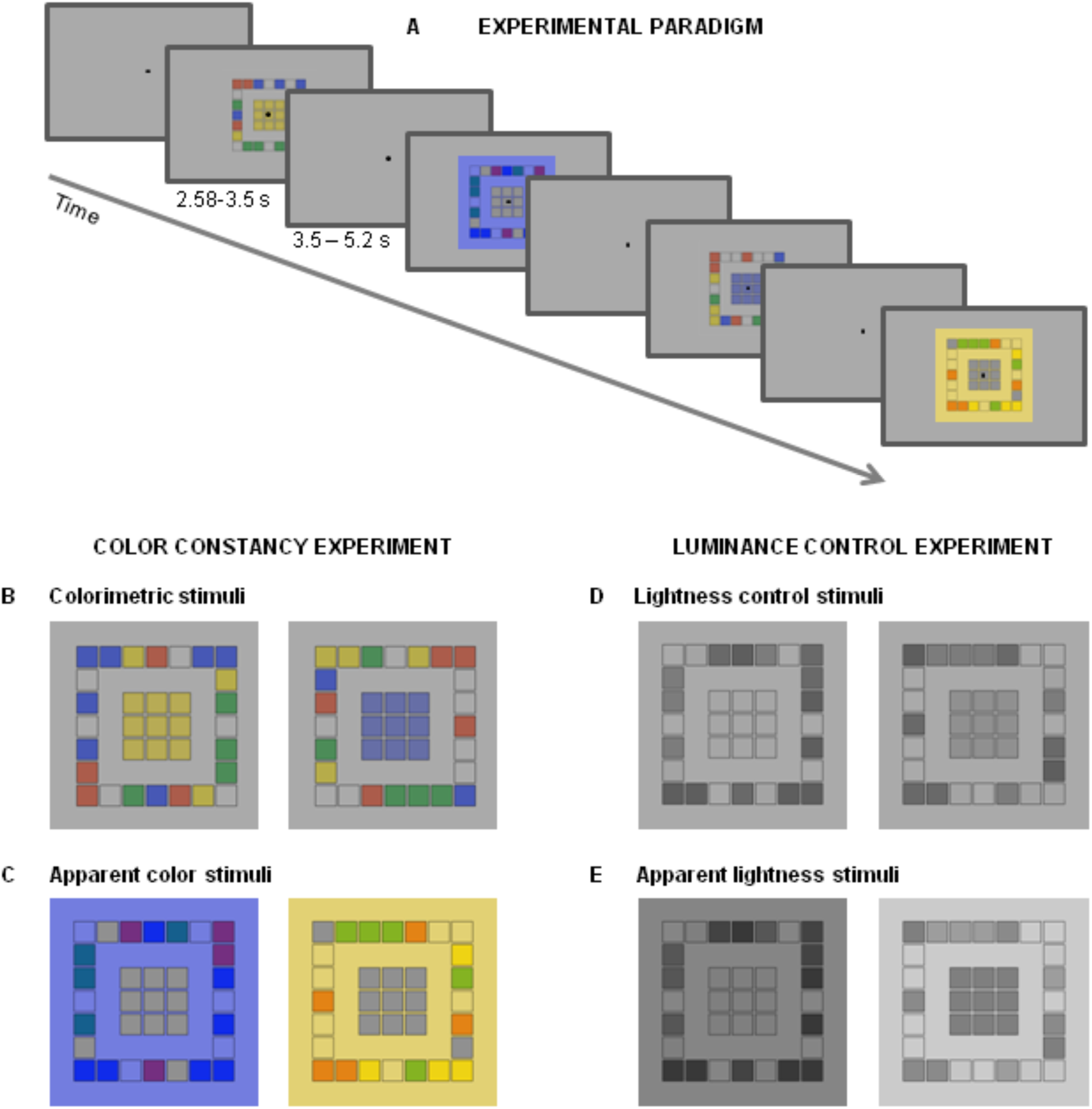
Experimental design. A) Experimental paradigm. Stimuli were presented in a random order for 2.5-3.8 seconds depending on oscillation speed) with a jittered interval of 3.5-5.2 seconds (2-3 TRs). Participants pressed a button whenever the fixation point flickered. Scanner runs contained apparent and colorimetric colors intermixed (day 1), or grayscale control stimuli intermixed (day 2). B) Colorimetric color stimuli. Stimuli were based on the cube by Lotto & Purves (2004), but adjusted for use in the MRI. In the colorimetric color condition, the center patches were yellow or blue, under a neutral illuminant. C) In the apparent color condition, the center patches were gray, surrounded by either a blue or a yellow illuminant, which caused the center patches to appear yellow or blue resp. D) Lightness control stimuli: grayscale versions of the colorimetric color conditions. E) Apparent lightness stimuli, which were grayscale versions of the apparent color stimuli. For each stimulus in B-E), there were six exemplars that randomly varied in the colorization of the surrounding patches.

In order to optimize the stimuli for the fMRI, we created 2D versions of the cube and increased the number of neutral gray center patches from 1 to nine (∼five degrees of visual angle), in order to create a stimulus that would activate a larger part of visual cortex, and therefore lead to more voxels we could use for decoding. We retained the small patch size to optimize the induction effect. Finally, we increased the gap between the center patches and the surround (Figure 1C) to separate the context from the region of interest. Although with the small patches, some surround might ‘leak’ into our area of interest, this should not affect the soundness of the effects: as the surround color (e.g. blue) is the opposite of the apparent, to-be-decoded color (e.g. yellow), classifying it as the surround (e.g. blue) would be incorrect, leading to lower classification accuracy. We made six different versions of each stimulus (900 x 900 pixels), in which the surround patches were randomly assigned one of the five colors from the original cube demo.

The presented stimuli were dynamically modulated to appear more or less colorful, in order to enhance the fMRI response. They gradually oscillated between the yellowish illuminant and an almost neutral illuminant at 40% of the yellowish illumination, or between the bluish illuminant and an almost neutral illuminant at 40% of the bluish illumination. We did not go below 40% in order to minimize the color aftereffects. To create neutrally illuminated versions of the scenes, we determined the point halfway between the RGB values for the yellowish and the bluish illumination for the neutral illumination, which appeared gray. The intermediate images were obtained by linearly combining the neutral and blue or yellow illuminated images at different proportions, ranging from 100% of the illuminated stimulus to 40% of the illuminated stimulus. The stimuli oscillated at 1-1.4 Hz and went through two full cycles. As a result, the illumination seemed to fade in and out, thereby causing the central patches’ appearance to oscillate between the color opponent to the illumination color and gray, even though the RGB-values of the center patches stayed constant throughout the whole time.

#### 2.2.2 Colorimetric Color Condition

The stimuli for the colorimetric condition (Figure 1B) largely resembled those for the apparent color condition. However, here we used a neutral illuminant and filled the central patches with a blue or yellow that perceptually matched the apparent blue and yellow of the apparent color condition (Figure 1C). To mimic the oscillation of the apparent color condition, the colorimetric stimuli then were made to oscillate between a version with neutral gray patches in the middle of the display, and a version with the blue or yellow patches in the middle of the display, both under a neutral simulated illuminant. The oscillation and stimulus-presentation durations matched those of the apparent color condition. These colorimetric stimuli were used to obtain the neural representation of “blue” and “yellow”, which we could subsequently use to test the representation of apparent blue and yellow.

#### 2.2.3 Lightness Control Stimuli

The lightness control stimuli were created from the colorimetric and apparent color stimuli by converting them to grayscale. First, the coordinates of each pixel were converted from RGB to DKL space. Then we set S-cone as well as L- and M-cone values to zero and converted all coordinates back to RGB. The grayscale version of the colorimetric stimuli contained a lightness difference (lightness stimuli, Figure 1D). The resulting grayscale stimuli of the apparent colors contained a lightness illusion (apparent lightness stimuli, Figure 1E): under the gray version of the blue illuminant, the center patches appeared lighter than under the gray version of the yellow illuminant. The control stimuli were used to test how much of the illusion and of the apparent color effect could be ascribed to lightness differences. The grayscale control stimuli also oscillated between background gray and the gray scale of the center patches (colorimetric color condition) or illuminants (apparent color condition).

### 2.3 Stimulus presentation and tasks

#### 2.3.1 Apparent Color & Colorimetric Color

The colorimetric and apparent color stimuli were presented intermixed in random order using an event-related design (Figure 1A). Each run contained 24 stimuli, with six presentations of each stimulus type (colorimetric blue, colorimetric yellow, apparent blue, apparent yellow) per run. Stimuli were separated by a jittered gap of 2-3 TRs (3.48 – 5.24 seconds). Four times per run, randomly interspersed with the colorimetric stimuli, a baseline condition was presented that contained a black outline of the patches on the neutral gray background. There were 12 runs in total, each of which lasted 3.7 minutes. Participants performed an attention-demanding task at the center of gaze. They had to fixate a central black fixation point and press a button whenever it seemed to flicker (i.e. it disappeared for one frame).

#### 2.3.2 Grayscale control

The grayscale control experiment was similar to the color induction experiment, except that grayscale versions of the images were shown. There were 24 trials per run, again randomly interspersed with the four baseline gray patch-outlines that were also presented in the colorimetric condition. Participants again had to fixate the central black fixation dot and press a button whenever it appeared to flicker.

#### 2.3.3 Functional localizers

In the functional localizers for the center and surround parts of the visual field, participants observed flickering gray checkerboard patches. In the center-localizer the checkerboards spatially matched the nine center patches of the induction experiment; in the surround-localizer the checkerboards matched the full surround of the center patches including the illuminated portion of the screen, but excluding the nine center patches. In order to enhance the BOLD response in the retinotopically relevant cortex the checkerboards flickered on and off, at a frequency of 4 Hz for 5.19 seconds (3 TRs), after which there was a blank period of 4.3 seconds. There were 30 blocks of the 5.19 s checkerboard presentation for the center-localizer and 30 blocks for the surround localizer. Participants fixated a red fixation dot in the center of the screen and pressed a button whenever it flickered.

#### 2.3.4 Retinotopy

We performed standard retinotopic mapping procedures to identify the borders between cortical areas V1, V2, V3 and V4 (S. A. Engel et al., 1997; Wandell et al., 2007). Retinotopy stimuli consisted of a rotating wedge and an expanding circle, which were presented in separate runs. The wedge and circle were filled with a high contrast, black and white checkerboard pattern that changed phase with a frequency of 4 Hz. The wedge stimulus completed 13 full cycles and the circle completed 10 full cycles. Each cycle took 33.75 seconds to complete. Participants had to fixate a red fixation dot in the center of the screen and were asked to press a button whenever they noticed a de-saturation of the fixation dot.

### 2.4 Technical setup

#### 2.4.1 Scan parameters

Data was collected with a SIEMENS Prisma 3 Tesla MR with a 64 channel head neck coil (Siemens AG, Healthcare, Erlangen, Germany). An anatomical scan was collected in 176 T1-weighted sagital images by means of an MP-RAGE sequence (voxel size 0.9 × 0.9 × 0.9 mm). A field map scan was acquired to measure the inhomogeneity of the magnetic field. All functional data were collected using a T2*-weighted gradient-echo planar imaging (EPI) sequence, with 25 slices (partial volume), acquired in descending order (slice thickness 3 mm; 0.8 mm gap; TR = 1.72 s; TE = 30 ms; flip angle 67°; field of view 220 × 220 mm; matrix size 74 × 74; voxel size 3 × 3 × 3 mm). Stimulus presentation and scanning parameters for the localizers were identical to those of the main experiment.

#### 2.4.2 Stimulus presentation

Stimuli were presented with a 1920 x 1080mm BOLDscreen 32 MRI-safe display (Cambridge Research Systems Ltd, Rochester, UK) behind the scanner bore. The visual stimulation could be seen by means of a double mirror attached to the head coil (visual field 28° in horizontal). Our stimuli encompassed approximately 14° x 14° degrees of the screen. We used Presentation software (Version 16, Neurobehavioral Systems TM, Albany, CA, USA) for stimulus presentation and response registration.

### 2.5 Analysis

#### 2.5.1 Preprocessing

Preprocessing was performed using MATLAB (Mathworks, Natick, MA) and SPM12 (Statistical Parametric Mapping, http://www.fil.ion.ucl.ac.uk/spm/). For each scanning session and run, functional scans were unwarped and realigned using the voxel displacement maps based on the measured fieldmaps. Then, the mean image of each run was co-registered with the first functional scan of the first scanning session, taking all the other images of that run along. The anatomical scan was also co-registered with the first functional scan. Since the multivariate analyses were performed in participant native space, no further preprocessing was performed before decoding. For the univariate analyses we subsequently normalized the functional and anatomical data to the Montreal Neurological Institute (MNI) space (using the SPM template) and smoothed the functional data with an 8mm FWHM kernel. For later visualization we created a mean anatomical image in MNI space by averaging the normalized anatomical images of our 19 participants.

#### 2.5.2 Univariate Model – Activation based analysis

To investigate which brain regions were activated by our colorimetric and apparent colors, we performed the first level analysis in the framework of the General Linear Model (GLM). We modeled the presentation of the colored stimuli using four separate regressors: colorimetric yellow, colorimetric blue, apparent yellow and apparent blue. For each regressor, we also included the temporal and dispersion derivatives in the model. Furthermore, we modeled the six head movement parameters (motion in the x-, y- and z-directions and pitch, roll, yaw) as regressors of no interest. To obtain sufficient baseline activity, we did *not* model the gray patch outlines that were presented four times in each run but included them in the implicit baseline. We then created the following contrasts: [colorimetric & apparent > baseline], which contained information about the presentation of *colored* stimuli, and [colorimetric > apparent], [apparent > colorimetric] to obtain activation purely to stimuli under neutral illumination or colored illumination. Finally, we created contrasts for yellow and blue stimuli versus baseline ([colorimetric yellow > baseline], [apparent yellow > baseline], [colorimetric blue > baseline], and [apparent blue > baseline]). The different contrasts at the first level were then input to a one-sample *t*-test at the second level and thresholded at 0.05 *FWE* at the voxel level, with no minimal cluster extent. For visualization, the results were overlaid on the averaged anatomical template.

#### 2.5.3 Retinotopic analysis, ROIs and voxel selection

Voxel selection consisted of two steps: first the delineation of visual regions V1-V4 using standard retinotopic mapping procedures (S. A. Engel et al., 1997), and secondly the selection of voxels within these regions that were most responsive to the center patches.

The retinotopic data were unwarped and realigned, co-registered, and then smoothed with 5 mm FWHM. The anatomical scan was inflated to reconstruct the cortical surface using Freesurfer (Martinos Center for Biomedical Imaging, Boston, MA, http://surfer.nmr.mgh.harvard.edu). We determined retinotopic areas V1, V2, V3 and V4 by means of a phase-encoded retinotopic mapping approach. A Fast Fourier Transformation was applied to each voxel’s time series to identify activation patterns that corresponded to the frequencies of the wedge and ring stimuli. The phase lags (i.e. the resulting polar angle and eccentricity maps) were then overlaid onto the reconstructed, inflated cortical surface. We delineated the borders of V1, V2 and V3 at reversals in the polar angle map. V4 was defined according to a recent review by Winawer & Witthoft (2015): the V3/V4 boundary was identified by a reversal in the polar angle representation, and the border towards VO1 was identified by an eccentricity reversal. The resulting masks served as Regions-Of-Interest (ROIs) in the further analyses.

Within each ROI, we determined the voxels that were most responsive to the center patches using the center-localizer and the surround-localizer. We created a GLM in which we modeled the visual stimulation with the checkerboards for center and surround separately. We also modeled the temporal and dispersion derivatives of the visual stimulation, as well as the six head movement parameters. We obtained contrasts for center stimulation [center > baseline], surround stimulation [surround > baseline], the unique contributions of center and surround ([center > surround] and [surround > center]), and their combination [center & surround]. Within each retinotopic ROI we then included only those voxels that were responsive to the center localizer (*t-*values > 0). Since there was a large overlap in activation for the center and the surround localizers, as a control, we also created a set of restricted ROIs, which contained only voxels that were sensitive to the center localizer (*t*-values > 0), but *not* to the surround localizer (*t*-values ≤ 0). We limited the number of included voxels to 100 per ROI. If there were more voxels that conformed to the criteria, we selected those with the highest *t-*vales for the center stimulation. For the restrictive ROIs, the number of available voxels was generally much smaller than 100 voxels, strongly limiting the input to the classifier (Figure S3C).

Finally, we created an additional ROI covering the Rolandic Operculum, as this region was consistently activated for both colorimetric and apparent stimuli. We used Marsbar (Brett et al., 2002) to create a sphere with a radius of 12 mm around the voxel with peak activation for the group contrast [colorimetric & apparent > baseline]. The resulting sphere was then converted back to the participants’ native space and included as an ROI in the analysis.

#### 2.5.4 Decoding analyses and overall cross-decoding logic

All decoding analyses were performed using The Decoding Toolbox (TDT (Hebart et al., 2015), Support Vector Machines (SVM) and custom MATLAB scripts. To test whether color representation could be decoded in the brain, we first trained and tested the classifier on colorimetric blue and yellow. Furthermore, we assessed whether there was a difference between apparent blue and yellow by training and testing an SVM on the apparent stimuli. Subsequently, we tested for color constancy: a color constant observer should perceive the gray center patches as yellow under blue illumination and blue under yellow illumination. A color constant representation therefore should be similar for colorimetric yellow and apparent yellow, or for colorimetric blue and apparent blue. To test this similarity, we trained an SVM to discriminate colorimetric blue and yellow, but tested whether it could also discriminate the apparent blue and yellow. Above chance decoding in this case indicates that the neutral patches are represented as yellow or blue, and hence that the color-constancy was achieved.

Since the blue and yellow stimuli contained a lightness difference on top of the hue difference, we performed a control experiment to test for the role of lightness and lightness constancy using the grayscale versions of the stimuli. We performed the exact same analyses as for the color experiment: lightness decoding (cross-validated), apparent lightness decoding (cross-validated) and a lightness constancy decoding, where we trained on the stimuli containing a lightness difference and tested on stimuli with an apparent lightness effect.

Finally, to test how much of the color constancy effect could be explained by lightness, we performed two more analyses. First, to test whether lightness plays a role in apparent yellow and blue, we trained the classifier to discriminate between stimuli of different lightness, that is, the gray versions of the colorimetric blue and yellow stimuli, and tested on apparent yellow and blue. Above chance decoding in this case would suggest that participants also achieve lightness constancy in their neural representations. Finally, we tested whether the SVM trained on colorimetric yellow and blue used the lightness difference in its discrimination by testing it on the stimuli with an apparent lightness difference. Above chance decoding in this case indicates that lightness is taken into account by the color-constant classification as well.

#### 2.5.5 Multivariate analysis: trial-wise classification

Activity in response to the relevant stimulus (colorimetric blue or yellow, or apparent blue or yellow) was calculated for each individual trial using a method based on the work by Mumford and colleagues (Mumford et al., 2014, 2012). Using single-trial estimates has the advantage that there are more exemplars available for training, which should allow the classifier to do better at generalization (Pereira et al., 2009). This is required for testing color-constant representations.

For each trial, we created a new GLM, where we modeled the respective trial in a single regressor and all other trials together in a second regressor. We also included the movement parameters as nuisance regressors. We repeated this for each the trial and obtained contrasts for that trial versus the implicit baseline. The resulting *t*-maps, which contained information about the specific activity in that one trial, were then input to the decoder.

We used a cross-validation approach when training and testing the classifier within colorimetric or within apparent colors. Each iteration we left out one run and trained on the remaining eleven runs. We repeated this twelve times, once for each run. The resulting accuracy was obtained by averaging the outputs from all twelve iterations. For generalization, we trained on all trials with colorimetric stimuli from all runs, and tested on all trials with apparent stimuli from all twelve runs as well.

#### 2.5.6 Multivariate analysis: run-wise classification

Since trial-wise estimates might lead to somewhat noisier estimates than run-wise estimates (Mumford et al., 2012), as a control analysis, we also calculated the mean response to each color per scanner run and used that as input for the decoder. Activity to the different colors per run was estimated using a GLM that contained regressors for colorimetric yellow, colorimetric blue, apparent yellow and apparent blue. For each regressor, we also included the first and second derivative in the model. Furthermore, we modeled the movement parameters as regressors of no interest. This GLM resembled the GLM used for the activation analysis, except that in this case the input was functional data in participant space, rather than MNI space. We then created contrasts for each color against the implicit baseline. The resulting *t*-maps were input for the classifier, as they should be more stable than the betas (Misaki et al., 2010).

We again used a leave-one-run out cross-validation approach when training and testing the classifier with colorimetric or with apparent colors. For each of twelve iterations the four beta’s of one run (colorimetric yellow and blue; apparent yellow and blue) were left out when training the classifier, and used for testing the classifier. The resulting accuracy was obtained by averaging the outputs from the twelve iterations. For generalization, we trained on colorimetric stimuli from all runs, and tested on apparent stimuli from all twelve runs as well.

#### 2.5.7 Permutation testing and statistical assessment

For decoding accuracies, the shape of the null distribution cannot be assumed to be normal and centered at 50%, therefore we used permutation tests to determine the significance of the actual decoding accuracies (Stelzer et al., 2013). For each participant, ROI, and decoding condition, we ran 1000 permutations, where we randomly shuffled the labels of the training and the testing data, following the procedures of The Decoding Toolbox (Hebart et al., 2015). Subsequently, we randomly drew a value from each participant’s individual permutation distribution and averaged the values from the 18 participants. We repeated this procedure 10000 times, each time randomly drawing a new value (with replacement) from each participant’s 1000 permutation results. The result was a null distribution of average decoding accuracies with 10000 values for each ROI and hypothesis.

In order to test whether the observed mean accuracies where inside or outside the range of what we could observe by chance, given that there was no information in the brain activity patterns, we checked the accuracy for each ROI and condition versus the distribution of mean accuracies from the permutations. We calculated how many of those permuted observations were *as high as, or higher than,* the actual observed mean accuracy. We then converted these to percentages, which directly reflected a *p-*value. We corrected for multiple comparisons over ROIs using the Holm-Bonferroni correction. Finally, we tested whether color-constancy decoding related to the decoding of colorimetric colors using Spearman rank correlations over participants.

#### 2.5.8 Searchlight analysis

To test whether any regions beyond V4 showed color constancy we employed searchlight analyses. We performed searchlight decoding for each combination of stimuli included in the ROI based analyses. We again used The Decoding Toolbox (TDT) to iteratively search through the brain using a searchlight sphere with a radius of 5 voxels. We masked the searchlight using the gray matter mask, as to include only the information of those voxels that were actually part of the brain. The resulting accuracy maps where then normalized to MNI space and smoothed with a 6 mm FWHM kernel. To asses which regions showed significant information in at least some of our participants, we submitted these normalized accuracy maps to a standard second level *t-*test in SPM.

## 3. Results

### 3.1 Univariate analyses: Overall activation in response to the (colored) stimuli

Colorimetric (neutral illuminant) and apparent (colorful illuminant) stimuli strongly activated a large cluster in the occipital lobe and the cuneus (Figure 2 and full results in Table S2), including V1, V2, V3, hV4, V01 and L01, as well as parts of V02, V3b and putative V4α (pV4α; Bannert and Bartels, 2017)). They also activated a small region in the frontal cortex, the Rolandic Operculum (MNI peak voxel: -42, -7, 17; 4 voxels, *t* = 7.45, *p_FWEcorr_* = 0.01, corrected for multiple comparisons using *Family-Wise Error correction* (*FWE-correction*) at a threshold of *p* < 0.05 at the voxel level).

**Figure 2:**
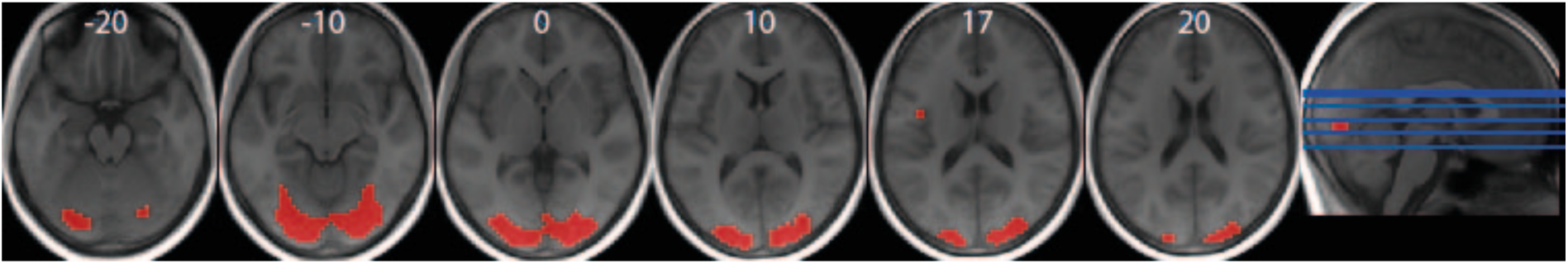
Univariate responses to colored and apparent stimuli on the average normalized brain. Slice numbers are indicated in white and their location is also presented on the sagital view of the brain. For visualization, activation is thresholded at *t* = 6.29. Peak activation was at MNI coordinates [18 -94 11], with sub peaks at [27 -91 -1] and [-27 -91 5], 1492 voxels, *t* = 25.45, *p_FWEcorr_* < 0.001. Full results can be found in Table S1.

Both blue and yellow colorimetric and apparent stimuli activated these same (early) visual regions (Figure S1B and S1C and Table S3). Larger activation for colorimetric stimuli than for apparent stimuli could be observed only in a small cluster (11 voxels) in V3d, while apparent colors led to larger activation in more anterior parts of the visual cortex and the fusiform gyrus (Figure S1D, Table S4). The center and surround localizers both activated large portions of early visual cortex (Figure S1A).

### 3.2 Multivariate Model – representation based analysis

#### 3.2.1 Decoding of colorimetric colors and color-constant colors in V1-V4

Colorimetric yellow and blue could be discriminated in all four visual regions V1, V2, V3 and V4 (V1: 53.95%, *p_corr_* < 0.0001; V2: 56.06%, *p_corr_* < 0.0001; V3: 55.90%, *p_corr_* < 0.0001; V4: 55.18%, *p_corr_* < 0.0001, permutation tests, see 2.5.7). This is in line with previous studies that have suggested color information is prominently processed throughout early visual cortex (Conway et al., 2007; S. Engel et al., 1997; Nasr et al., 2016; Tootell et al., 2004). Training and testing on apparent colors also led to significant decoding accuracies in all four ROIs V1-V4 (Figure S2A V1: 58.89%, p_corr_ = 0; V2: 59.59%, p_corr_ = 0; V3: 60.23%, p_corr_ = 0; V4: 58.32%, p_corr_ = 0; permutation tests). To test for color-constant representations in early visual regions, we trained the classifier on the colorimetric colors and tested on the apparent colors, because a color-constant representation should resemble that of colorimetric yellow and blue. Discrimination of apparent yellow and blue was possible in V1, V3, and V4 (Figure 3A), with a trend towards significance in V2 (V1: 53.12%, *p_corr_* < 0.0001; V2: 51.43%, *p_corr_* = 0.06; V3: 54.47%, *p_corr_* < 0.0001; V4: 51.95%, *p_corr_* = 0.019; permutation tests). When we repeated the decoding analyses using run-wise regressors we found significant decoding in V1 and V2, and a trend in V3, suggesting that for all these ROIs, decoding accuracies are consistent (Figure S3B; V1: 54.42%, *p_corr_* = 0.02; V2: 56.46%, *p_corr_* = 0.003; V3: 53.20%, *p_corr_* = 0.072; V4: 51.12%, *p_corr_* = 0.32; permutation tests). Although it cannot be excluded that certain voxels responded to the surround illumination as well, the fact that the surround color was the opposite of the apparent (and to-be-decoded) color should have lowered the decoding accuracy, rather than enhanced it. However, to control for this we repeated the analyses using the ROIs restricted to voxels not responsive to the surround localizer and found above chance decoding in V2 and V3 (Figure S3A; V1: 50.78%, *p_corr_* = 0.21; V2: 53.51%, *p_corr_* < 0.0001; V3: 53.61%, *p_corr_* < 0.0001; V4: 50.20%, *p_corr_* = 0.039; permutation tests).

**Figure 3:**
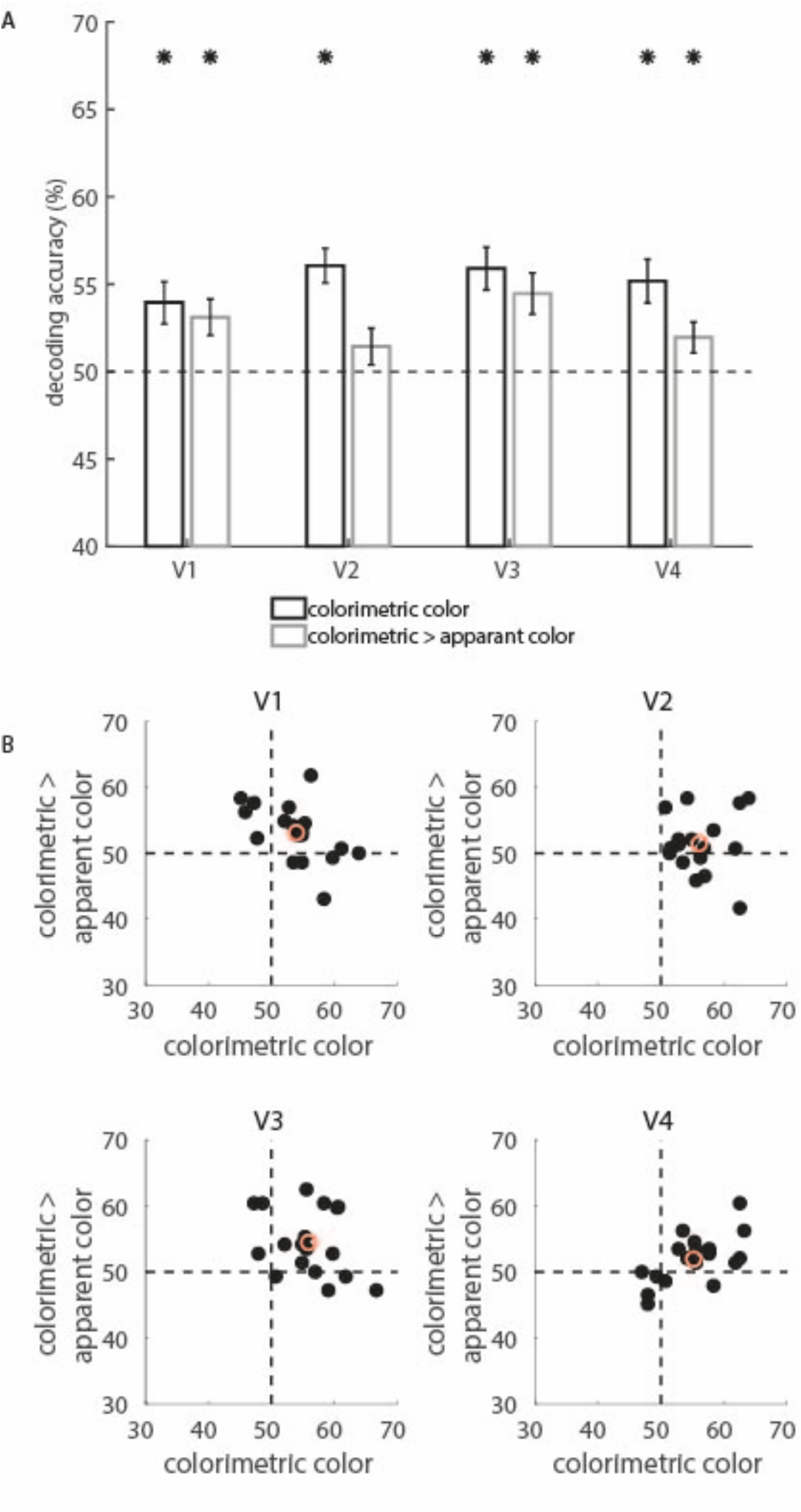
Decoding of apparent and colorimetric stimuli. A) Decoding accuracies (%) for colorimetric colors (black) and apparent colors (gray) in V1-V4. Color constant colors were obtained by training the classifier on the colorimetric colors, while testing on the apparent colors. Bars indicate average decoding accuracy for each condition and ROI. Black dashed line indicates the theoretical chance level (50% accuracy). Black asterisks indicate *p* < 0.05 corrected for multiple comparisons; obtained from the permutation tests. B) Relationship between decoding accuracies for colorimetric stimuli and for generalization between colorimetric and apparent colors, for V1-V4. Black dots indicate the accuracy of each individual participant. The red circle indicates the mean decoding accuracy. Black dashed lines indicate chance level.

Finally, even though there was some variation between participants in how well we could decode colorimetric colors and color-constant colors, better colorimetric decoding did not lead to better color-constant decoding (Figure 3B, all *p* > 0.27 uncorrected).

#### 3.2.2 Decoding lightness and generalizing to lightness constancy in the gray control stimuli

To test how much of the observed effects for color constancy could be explained by lightness constancy, we performed a control experiment in which participants observed only grayscale versions of the colorimetric and apparent color stimuli: the lightness and apparent lightness stimuli. Decoding lightness was possible in V1-V4 (Figure 4A; V1: 57.52%, *p_corr_* < 0.0001; V2: 58.86%, *p_corr_* < 0.0001; V3: 54.83%, *p_corr_* < 0.0001; V4: 53.95%, *p_corr_* < 0.0001; permutation tests). The lightness difference also generalized to apparent lightness: in V1-V4 lightness was represented in a constant fashion where a classifier trained on actual lightness could be used to discriminate apparent lightness (V1: 54.65%, *p_corr_* < 0.0001; V2: 56.11%, *p_corr_* < 0.0001; V3: 53.33%, *p_corr_* = 0.002; V4: 52.15%, *p_corr_* = 0.022; permutation tests). Training and testing within apparent lightness, finally, was possible in all four areas (Figure S2B). Similarly to the apparent colors, this could be an effect of a perceptual difference in the surround. Better lightness decoding related weakly to better discrimination of lightness-constancy in V1 (*r* = 0.48, *p* = 0.09, Figure S2C).

**Figure 4:**
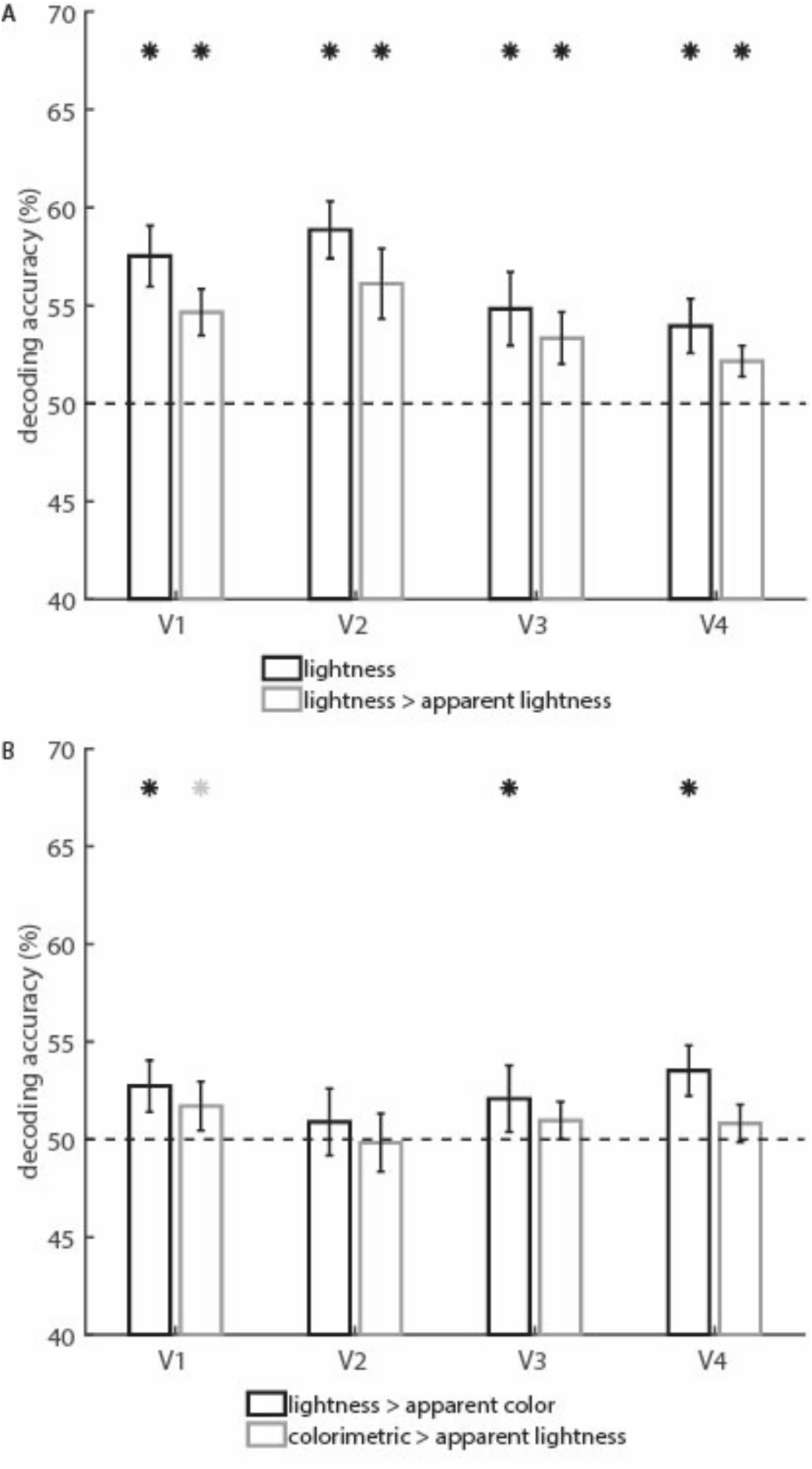
Decoding accuracies for lightness control stimuli in V1-V4. A) Decoding accuracies for lightness stimuli (black) and lightness constancy (gray) in V1-V4. Lightness constancy was obtained by training the classifier on the gray version of colorimetric colors, while testing on the gray version of the apparent colors. Bars indicate average decoding accuracy for each condition and ROI. B) the effect of lightness n apparent color (black) obtained by training on lightness stimuli and testing on apparent color and the effect of lightness in the color classifier (gray) by training on colorimetric color stimuli and testing on apparent lightness. Bars indicate average decoding accuracies over 14 participants. Black dashed line indicates theoretical chance level (50%) and black asterisks indicate p < 0.05 obtained from the permutation tests, corrected for multiple comparisons; gray asterisks indicate *p* < 0.05 uncorrected.

#### 3.2.3 Testing the role of lightness in the apparent color

To test whether the lightness difference could partly explain the decoding of the apparent colors, we trained classifiers to discriminate between the lightness control stimuli and tested them on the colored stimuli, and vice versa. The classifier trained on the lightness difference was able to discriminate between apparent blue and yellow responses, in V1, V3 and V4 (Figure 4B; V1: 52.73%, *p_corr_* = 0.011; V2: 50.89%, *p_corr_* = 0.19; V3: 52.08%, *p_corr_* = 0.036; V4: 53.52%, *p_corr_* = 0.0007; permutation tests). However, the decoding of this classifier was weaker than for the classifier trained on colorimetric stimuli. The classifier trained on the difference in response to colorimetric yellow and blue was not able to discriminate apparent lightness, although it was significant in V1 at an uncorrected level (Figure 4B; V1: 51.71%, *p_corr_* = 0.19 (uncorrected *p* = 0.048); V2: 49.83%, *p_corr_* = 0.57; V3: 50.96%, *p_corr_* = 0.52; V4: 50.81%, *p_corr_* = 0.52; permutation tests). Finally, there was no correlation between how well lightness explained the color-constancy and how much the classifier based on colorimetric yellow and blue relied on lightness differences. Only V2 showed a trend towards a significant correlation (Spearman’s *r =* 0.51, *p* = 0.065 uncorrected).

#### 3.2.4 Decoding in the Rolandic operculum

Since the Rolandic operculum showed activation for colorimetric and apparent color stimuli, we created a sphere around the peak voxel and used it as an ROI in all the decoding analyses. It was possible to discriminate between apparent colors (52.20%, *p*_corr_ =0.025), and somewhat possible to discriminate between apparent lightness levels (52.01%, p_corr_ =0.065). It was not possible to discriminate between colorimetric (51.18%, *p_corr_* = 0.14), or lightness stimuli (51.45%, p_corr_ = 0.13), nor was it possible to generalize between any of the conditions (all accuracy < 51.09%; all *p* > 0.13; Figure S5).

### 3.3 Searchlight decoding of color-constancy

We performed searchlight analyses for all combinations of stimuli that were also in the ROI-based analyses, to explore whether there were any other regions that represented colors in a colorimetric or color-constant fashion. All group average decoding maps were first thresholded at *p* < 0.001, and then corrected for multiple comparisons using *FWE-correction* at the cluster level.

The searchlight indicated that it was possible to decode colorimetric yellow and blue in the vast majority of the occipital lobe, including visual areas V1-V4 (Figure 5 and Table 1). Furthermore, above chance decoding was found in the right Insula, as well as in the anterior and middle Cingulum and superior frontal medial gyrus. Decoding of apparent colors was also possible in large portions of the occipital lobe (Table 1), including the pV4α-area reported by Bannert & Bartels (Bannert and Bartels, 2017), as well as in one frontal area (MNI: [-54 -7 29], *t* = 4.59, *p_FWE-_*_corr_*=* 0.027), corresponding to the (pre-)central gyrus, and close to the Rolandic operculum. Finally, the searchlight trained on colorimetric colors and tested on apparent colors found color-constant representations only in an area in the right hemisphere (MNI: [30 -94 20], *t* = 5.11, *p_FWE-_*_corr_*=* 3.21*10^-6^), probably corresponding to dorsal V2 and V3.

**Figure 5:**
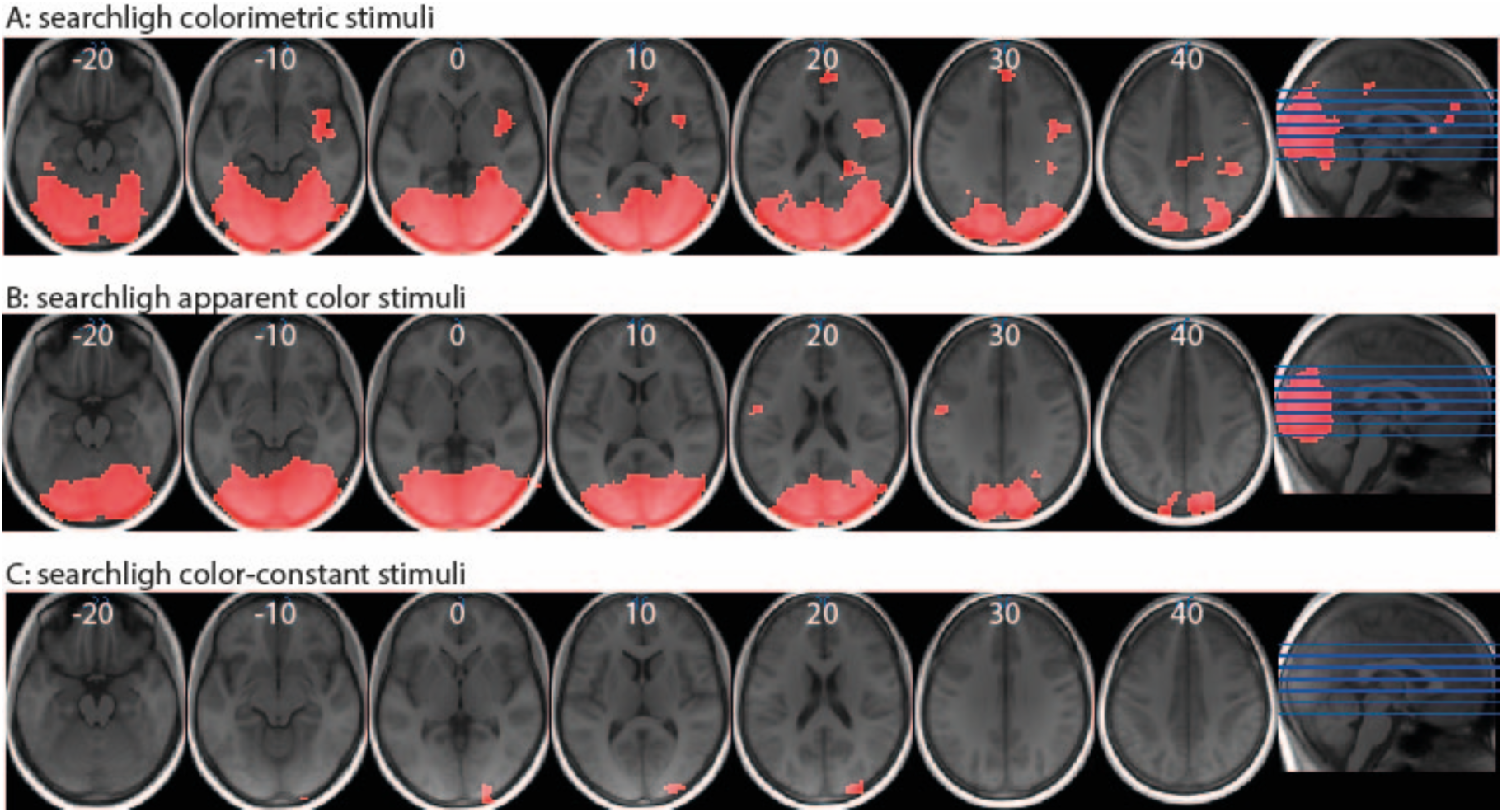
searchlight results. A) Above chance decoding for colorimetric stimuli, B) apparent color stimuli and C) color constant stimuli, overlaid on the average brain in MNI space. Slice numbers (MNI) are indicated in white, slice position is indicated on the sagital view of the brain (right-most figures). Activation maps are thresholded *p* < 0.001 uncorrected and then corrected at the cluster level (43, 169 and 48 voxels resp.), corresponding to a *t* = 3.65.

**Table 1:** Searchlight results, thresholded first at *p* < 0.001 uncorrected at the voxel level and subsequently corrected for multiple comparisons at the cluster level (43, 169 and 48 voxels resp.).

| Colorimetric colors | Cluster |  | Peak |  | Location |  |
| --- | --- | --- | --- | --- | --- | --- |
|  | p(FWE-corr) | Size (voxels) | T | X | Y | z |
| *Occipital cortex* | $p < 0.0001$ | 11050 | 12.11 | 33 | -79 | 8 |
|  |  |  | 11.19 | 24 | -70 | 8 |
|  |  |  | 11.14 | 24 | -85 | -10 |
| *Right insula* | $p < 0.0001$ | 439 | 6.13 | 33 | 2 | 23 |
| *Anterior cingulum, superior frontal middle gyrus* | 0.00036 | 101 | 5.05 | 3 | 44 | 29 |
| *Medial cingulum* | 0.045 | 43 | 4.43 | 15 | -28 | 35 |
| Color-constant colors | Cluster |  | Peak |  | Location |  |
|  | p(FWE-corr) | Size (voxels) | T | X | Y | Z |
| *Middle occipital gyrus, V2d, V3d* | $p < 0.0001$ | 169 | 5.11 | 30 | -94 | 20 |
| Apparent colors | cluster |  | Peak |  | Location |  |
|  | p(FWE-corr) | Size (voxels) | T | X | Y | Z |
| *Inferior occipital lobe* | $p < 0.0001$ | 8616 | 12.67 | 36 | -70 | -4 |
|  |  |  | 11.79 | 3 | -82 | 2 |
|  |  |  | 11.36 | 6 | -88 | -4 |
| *Left precentral gyrus, postcentral gyrus* | 0.027 | 48 | 4.59 | -54 | -7 | 29 |

## 4. Discussion

Early visual neural responses to apparent blue and yellow surfaces resemble the responses to colorimetric yellow and blue. Our classifier trained on BOLD activity patterns for colorimetric yellow and blue, was able to discriminate between activity patterns to apparent yellow and blue, induced by bluish and yellowish illuminants on neutral gray patches. This suggests appearance based neural coding of color in human early visual cortex. Decoding was significantly above chance in V1, V3 and V4, with a trend in V2 (Figure 3). However, there was no significant difference between the ROIs, suggesting that color-constant representations are present in all of them. While luminance and lightness constancy were also represented in these areas, lightness could not altogether explain the color induction findings (Figure 4), pointing to a genuine effect color-constant neural representation in the earliest cortical visual areas.

That most of occipital lobe, including V1-V4, is sensitive to hue has been shown before (e.g. Bannert & Bartels, 2013, 2018; Brouwer & Heeger, 2009, 2013; Goddard, Mannion, McDonald, Solomon, & Clifford, 2010; Hong & Tong, 2017; Kuriki et al., 2011; Vandenbroucke, Fahrenfort, Meuwese, Scholte, & Lamme, 2016). Our decoding results show that these responses in human observers are *appearance-based*, rather than based on the photometric receptor activations in the eye. Such early representation of color-constant stimuli in human V1 is in accord with the findings from monkey physiology (Wachtler et al., 2003, 2001). For human observers representations of apparent colors had been reported for V3 and V4 (e.g. Bannert and Bartels, 2017; Hong and Tong, 2017; Vandenbroucke et al., 2016), but such early appearance-based coding had not yet been shown aside from memory colors (Bannert and Bartels, 2013). A recent study did show illuminant-invariant representations surface colors pV4α and V1 with about the same magnitude of decoding (Bannert and Bartels, 2017), with the ability to discriminate surface colors in V1 driven by information about the illuminant. We confirm those findings in V1 and extend them with a color constant representation in V2, V3 and V4 (Figure 3A). In the searchlight analysis, area pV4α (Bannert and Bartels, 2017; Barbur and Spang, 2008) showed colorimetric and surface-color representations, but no generalization from colorimetric to apparent colors. Because of differences in stimulus design (e.g. 2D surfaces vs. rendered scenes) and classifier training approach (colorimetric stimuli under neutral illumination vs. colored stimuli under multiple different illuminants), the results from Bannert & Bartels (Bannert and Bartels, 2017) cannot be directly compared with the current study. Furthermore, it is important to note that a lack of decoding does not exclude a lack of representations; the BOLD response is a somewhat indirect measure of neural activity that can be measured more or less precisely depending on many factors, such as anatomy and spatial organization. In any case, both studies show (illuminant-invariant) surface representations across different early visual areas and specifically in V1.

V1 might be perfectly suited for further processing of color constancy, due to its ability to process local context. Sensitivity to edge contrast, one of the mechanisms thought to support color constancy (Foster, 2011), first emerges in primary visual cortex (V1), in so-called double-opponent cells (Conway, 2001; Conway and Livingstone, 2006; Johnson et al., 2001; Shapley and Hawken, 2002). A recent model (Mély and Serre, 2016) constrained by V1 anatomy and physiology, showed that a recurrent network of facilitatory and inhibitory input from the receptive fields in the near and far surround was able to predict exactly the hue shift that human observers perceived in their center-surround color stimuli (Klauke and Wachtler, 2015; Mély and Serre, 2016). This perceived hue shift also qualitatively matches the hue shift observed in V1 neurons under different illuminants (Wachtler et al., 2003).

Our findings of color-constant representations in V1, V2 and V3, as well as V4 ((Schein and Desimone, 1990; Zeki, 1983) agree with the idea that color constancy depends on processing at multiple sites in the color pathway (Barbur and Spang, 2008; Foster, 2011; Walsh, 1999). While V1 contains appearance-based representations and is able to calculate local contrast, it might not by itself be sufficient to obtain color constancy. First of all, several patients with color-constancy impairments have lesions around V4 (Clarke et al., 1998) or even in frontal cortex (Rüttiger et al., 1999). There are even reports of one patient with spared V1 and V2 and color awareness (Zeki et al., 1999), that nonetheless did not show constancy. Color constancy can also be relatively spared when color vision is gone (Cowey and Heywood, 1997). Secondly, response normalization at larger scales, such as assumed in the gray-world assumption, has to take place at later stages of the visual system, where receptive fields are correspondingly larger. These more global effects might propagate information back to early visual cortex and V1 might be the perfect place to integrate the bottom-up (retinal adaptation, local contrast) and top-down (illuminant discounting, memory color) information (Bullier, 2001; Roelfsema and de Lange, 2016) that can be used to calculate color-constancy. Such top-down inputs presumably also underlie the representations of memory colors in V1 (Bannert and Bartels, 2013). At the same time, most appearance effects for color representation seem to appear later than V1: color imagery representations were found in hV4 (Bannert and Bartels, 2018); color filling-in could be decoded from V3 and V4 (Hong and Tong, 2017); the representation of ambiguous color in typical objects was influenced by prior object-knowledge in V3 and V4 (Vandenbroucke et al., 2016), and representations of color were influenced by color-categories only in V4 (Brouwer and Heeger, 2013). The earlier, present throughout V1, V2 and V3, representation of color-constant colors underscores the idea that color-constancy is partly dependent on low level processes such as retinal adaptation and local contrast.

A candidate region for providing top-down input into visual cortex is the Rolandic Operculum (RO). We unexpectedly found this region to be activated during perception of colorimetric and apparent colors. The RO is located close to cognitive control regions (De Baene et al., 2012; Staddon, 2005) and has been implicated in articulation and (mental) verbalization (Tonkonogy and Goodglass, 1981; Zarnhofer et al., 2012). Possibly, participants might have verbalized the names of the apparent colors. However, the Rolandic Operculum also overlaps with the location of lesions that showed impairments in color-constancy (Rüttiger et al., 1999). Apparent blue and yellow, as well as apparent lightness, but not colorimetric blue and yellow, could be somewhat discriminated in this region. Presumably, the receptive field sizes in this frontal region might have prevented distinction between center and surround. At the same time, this renders the Rolandic Operculum well suited for context or illuminant processing, which might then be fed back to early visual cortex. Future studies could explore the exact role of this region, as well that of a nearby region in the precentral gyrus found with the searchlight analyses, in color constancy.

Both lightness information and lightness constancy where represented in V1-V4, as expected from previous work (Boyaci et al., 2007; Ruff et al., 2018). Given that illumination changes invariably lead to changes in the lightness of some surfaces in a scene, lightness is expected to play a role in color constancy. We therefore tested whether lightness differences in our stimuli could explain the color-constancy decoding results. Lightness seemed to indeed play a role in the representation of color-constant colors: a classifier trained on lightness differences could also discriminate somewhat between apparent blue and yellow in V1, V3 and V4 (Figure 4C). However, the classifier trained on colorimetric blue and yellow was not able to discriminate between apparent lightness levels (Figure 4C), suggesting that this classifier uses hue information and the color-constant decoding results are thus not solely due to lightness. Of course, to ultimately test whether lightness plays a role, one could create stimuli that are iso-luminant, such as in the recent study by Bannert & Bartels (2017). Yet, illuminant changes in daily life typically also involve lightness differences, and these do provide strong cues for color-constancy (Golz and MacLeod, 2002). Thus, to fully understand color-constancy, it might be necessary to understand lightness constancy and the (potentially same) mechanisms underlying it, as well.

The decoding accuracies we observed where relatively weak in absolute terms (∼55%), for the colorimetric, color-constant and lightness stimuli. First of all, since in the current study we are optimizing *interpretation* rather than *prediction* (Hebart and Baker, 2017), absolute decoding accuracies are less important than consistency over participants. Decoding accuracies are not standardized effect sizes, as they depend partly on the specific analyses procedures (Hebart and Baker, 2017). In the current work, the use of trial-wise regressors might have provided better generalization (Pereira et al., 2009), but also noisier estimates (Mumford et al., 2012).

Secondly, color constancy is not perfect perceptually: color constancy matches are often in the range of 40-60% (Foster, 2011; Witzel and Gegenfurtner, 2018), suggesting that the apparent and colorimetric colors might have not been perceived as equal, hampering decoding accuracy. Yet, the fact that decoding is possible shows that some similarity exists. The perceived extent of color constancy in our current stimuli might have been further limited by design and measurement conditions: the stimuli were relatively small compared to previous studies of color decoding (Bannert and Bartels, 2018, 2013; Brouwer and Heeger, 2013, 2009; Parkes et al., 2009), limiting the number of available voxels for decoding; the oscillations induced to prevent neural adaptation might have limited the perceptual effect; the central fixation task could have drawn participants attention away from the full scene; the stimuli were relatively artificial surfaces and illuminants created from simple 2D squares. Although it is still an open question which cues exactly play a role in color-constancy (Foster, 2011), real objects and scenes might provide more extensive clues on surface color. Yet the more realistic, rendered scenes in the study by Bannert & Bartels (2017) resulted in decoding accuracies within a similar range.

Finally, many voxels were responsive to both the center and the surround localizers, making it difficult to separate voxel activity for center and surround patches. Voxels selected to respond to the center patches, might have somewhat activated in response to the illuminants. The classifier might have used this ‘colorimetric representation in the surround’ for its discrimination between apparent yellow and blue. Because apparent blue is induced by the yellow illuminant, and vice versa, a classifier trained on colorimetric color that used surround information would classify apparent blue as yellow and vice versa. Thus, it would be effectively wrong mostly and hence lead to low decoding accuracies for generalization. Only for the classifier that was trained and tested on apparent colors, such use of the surround could lead to higher decoding accuracies: classifying blue as yellow and vice versa would be possible during both training and testing and hence lead to correct responses. Indeed, the highest decoding accuracies where found when training and testing within apparent colors. Therefore, with the current stimulus set-up, it is difficult to draw inferences about the strength of representation of the apparent colors per se; even though this higher accuracy could also be a genuine effect of lateral connections from the surround that send information to the foveal area. However, to reiterate, for the color-constant representations where the classifier was trained on colorimetric colors, this can mostly have harmed our classification. It will be a challenge for future studies to design stimuli that circumvent such practical limitations in the fMRI and optimize accuracy.

To conclude, we found representations of color-constant colors in early visual regions V1, V3 and V4, with a trend in V2. Although luminance differences might play a role in color-constancy, these representations could not be fully explained by lightness differences between the stimuli. Rather, these results support the idea that color constancy depends at several stages of processing in both visual areas and potentially higher order regions such as the Rolandic operculum.

## Acknowledgments

The authors would like to thank David Weiss for support in stimulus creation, Hanna Gertz and Axel Schafer for help with data collection and storage, and Benjamin de Haas and Sérgio Nascimento for useful feedback on design and analyses. The authors were supported by the Deutsche Forschungsgemeinschaft SFB/TRR 135 (grant number 222641018): Cardinal mechanisms of perception. Some of these results were first presented in abstract form at the 2016 VSS meeting (Baumgartner, Weiss & Gegenfurtner, 2016, Journal of Vision, Vol.16, 392).

## Supplemental results

### Activation to the center & surround localizers

Both the center and the surround localizers activated large portions of the occipital lobe (including Calcarine sulcus, inferior and middle occipital gyrus, and posterior parts of fusiform and lingual gyrus, Figure S1A). The center-localizer activated more posterior areas, as expected for stimulation around the fovea. Peak activations were found in the right hemisphere (MNI: [24 -91 -4], sub peaks at [24 -94 5] and [45 -70 -1], 747 voxels, *t* = 24.93, *p* = 0.000) and the left hemisphere (MNI: [-21 -88 -10], sub peaks at [-27 -94 2] and [-36 -85 -4], 548 voxels, *t* = 19.17, *p* = 0.000). These two clusters of activity included bilateral V1, V2, V3, hV4, V01 and L01.

The surround-localizer activity largely overlapped with center-localizer activity, but it extended into more anterior, peripherally-sensitive areas, which also included inferior and middle temporal gyrus. Peak activations were found in visual cortex (MNI: [12 -91 -1], with sub peaks at [27 -85 23] and [33 -82 5], 2374 voxels, *t* = 14.20, *p* = 0.000) which included bilateral functional areas V1, V2, V3, hV4, hMT, PHC1, IPS, V01, V02 and L02. Furthermore, there was a small cluster of activity in the hippocampus / thalamus (MNI: [21 -31 2], 3 voxels, *t* = 6.72, *p* = 0.030).

Although the stimuli for center and surround localizers did not overlap in visual space, there was a large overlap in regions that were activated by these stimuli. This overlap limited the possibility to select voxels that were exclusively responsive to center stimuli, as very few voxels showed such a profile. However, please not that even though selected voxels might have been also responsive to the surround, this should not affect the soundness of the decoding results: as the surround color is the opponent color to the apparent color, it should tip the classifier into the opposite direction, leading to lower classification accuracy.

### Activation to individual colorimetric and apparent stimuli

Colorimetric yellow and blue both activated early visual cortex (including V1-V4, V3A and parts of the fusiform gyrus), with yellow leading to somewhat more extensive activation (Figure S1B). Yellow MNI peak: [18 -94 8], sub peaks at [27 -91 -4] and [-27 -91 5], 1409 voxels, *t* = 26.33, *p* = 0.000. Blue MNI peak: [27 -91 -1], sub peak at [18 -94 8], 469 voxels, *t* = 20.67, *p* = 0.000 and [-24 -94 5], sub peaks at [-21 -85 -10] and [-18 -97 11], 538 voxels, *t* = 17.41, *p* = 0.000.

As for the colorimetric stimuli, apparent yellow led to slightly more extensive activation than apparent blue (Figure S1C); again including V1-V4, V3A and parts of fusiform gyrus. Yellow induction MNI peak: [18 -94 11], sub peaks at [27 -91 -1] and [-24 -88 -10], 1648 voxels, *t* = 18.63, *p = 0.000.*Blue induction MNI peak: [18 -94 11], sub peaks at [-18 -94 14] and [12 -91 2], 1368 voxels, *t* = 21.67, *p* = 0.000.

### Activation for colorimetric > apparent and activation for apparent > colorimetric

Apparent color stimuli activated a larger part of the brain, mostly encompassing the more anterior portion of early visual regions, including V1, V3v and V3d. Such peripheral activation specific to apparent colors could be explained by the greater part of the visual field taken up by the apparent stimuli, because of the colored illuminant surrounding the center patches. Colorimetric stimuli specifically activated one small cluster (11 voxels) in what is most likely dorsal V3.

**Figure S1:**
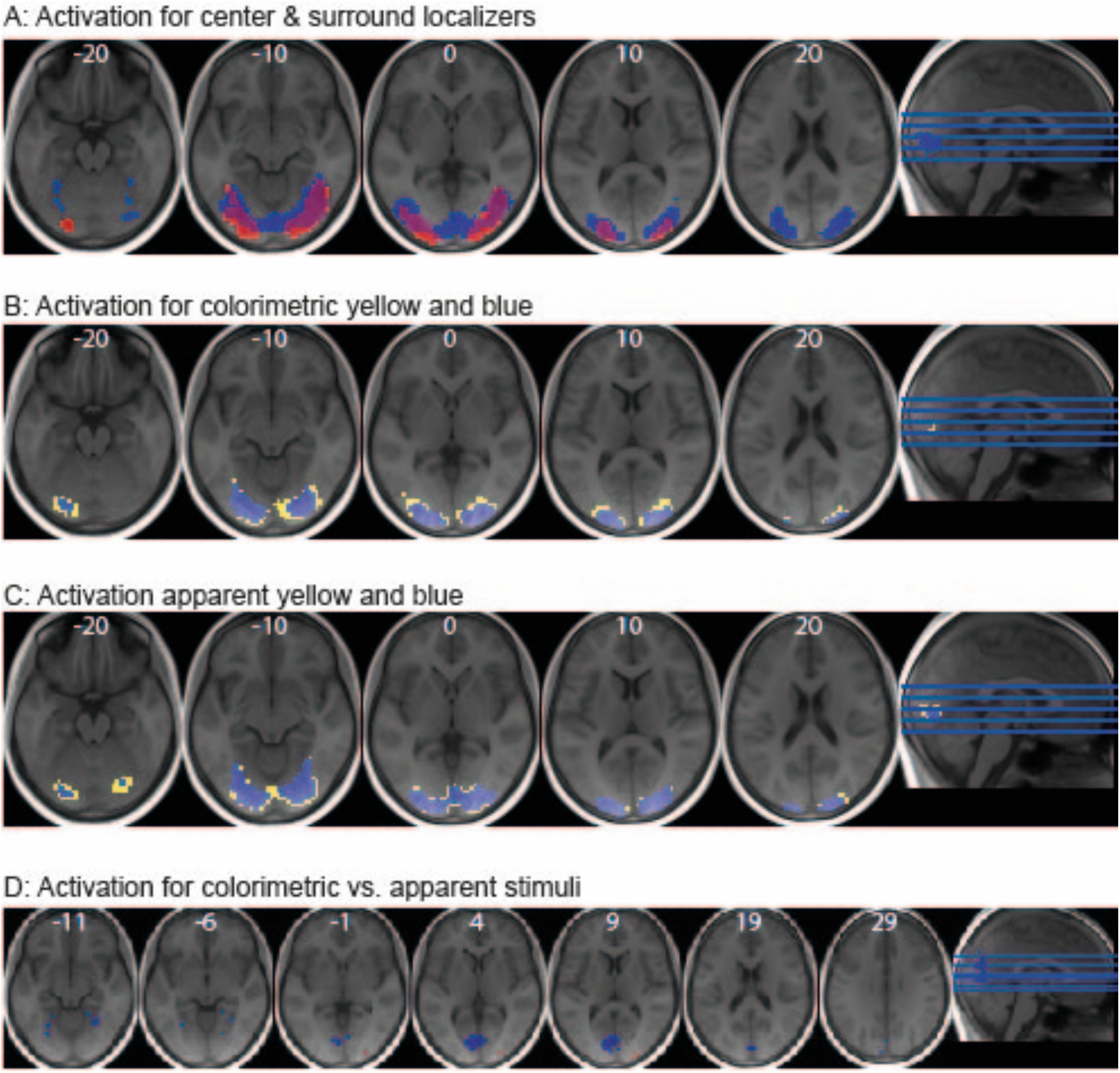
Univariate results. A) Voxel selection and activation to the center and surround. Group activations for the center localizer (red, *t* = 6.44) and the surround localizer (blue, *t* = 6.37) on the averaged brain (MNI space). Thresholded at *p* = 0.05 *FWE*-corrected, cluster extent = 0; in white the z-coordinates of each slice in MNI space and a visualization of the slice locations on a sagital slice (rightmost image). B) Activations for colorimetric yellow (yellow, t = 6.40) and blue (blue, t = 6.34), thresholded at *p* = 0.05 *FWE, cluster extent = 0;* as in A). C) Activations for apparent yellow (yellow, t = 6.33) and blue (blue, t = 6.29), thresholded at p = 0.05 FWE, cluster extent = 0; as in A). D) Activation that is larger to colorimetric stimuli than to apparent stimuli (red) and activation that is larger for apparent colors than for colorimetric stimuli (blue), thresholded at *p* = 0.05 *FWE, cluster extent = 0;* as in A).

**Table S1:** Activation for all stimuli (colorimetric and apparent color) compared to baseline. See also Figure 2.

| colorimetric & apparent > baseline | cluster |  | peak | location |  |  |
| --- | --- | --- | --- | --- | --- | --- |
| | $P_{(FWE-corr)}$ | Size (voxels) | T | x | y | z |
| *Cuneus, Calcarine sulcus, superior occipital gyrus, V2d* | $p < 0.0001$ | 1492 | 25.45 | 18 | -94 | 11 |
| *Inferior occipital gyrus, V3d* |  |  | 21.27 | 27 | -91 | -1 |
| *Middle occipital gyrus, V3d* |  |  | 19.00 | -27 | -91 | 5 |
| *Rolandic operculum* | 0.0099 | 4 | 7.45 | -42 | -7 | 17 |

**Table S2:** Activation for the center localizer (top) and the surround localizer (bottom) compared to baseline. The center localizer also activated V1, V3d, hV44 VO1, posterior fusiform gyrus, left middle/occipital gyrus, Calcarine sulcus and lingual gyrus. The surround localizer also activated V3, V2, V1, V01, V02, PHC1, hV4, V3a, IPS and L02. See also Figure S1A.

| Center Localizer | Cluster |  | peak | location |  |  |
| --- | --- | --- | --- | --- | --- | --- |
| | $P_{(FWE-corr)}$ | Size (voxels) | T | X | Y | Z |
| *Right inferior occipital gyrus, V2v* | $p < 0.0001$ | 747 | 24.93 | 24 | -91 | -4 |
| *Right middle occipital gyrus, v2d* |  |  | 18.30 | 24 | -94 | 5 |
| *Right middle temporal gyrus, LO1* |  |  | 12.49 | 45 | -70 | -1 |
| *Left inferior occipital gyrus, V3v* | $p < 0.0001$ | 548 | 19.17 | -21 | -88 | -10 |
| *Left middle occipital gyrus, LO1* |  |  | 16.03 | -27 | -94 | 2 |
| *Left inferior occipital gyrus, LO1* |  |  | 13.13 | -36 | -85 | -4 |

**Table S2: Activation for the center localizer (top) and the surround localizer (bottom) compared to baseline.**
| Surround Localizer | Cluster |  | Peak | location |  |  |
| --- | --- | --- | --- | --- | --- | --- |
| | $P_{(FWE-corr)}$ | Size (voxels) | T | X | Y | Z |
| *Right calcarine, L02* | $p < 0.0001$ | 2374 | 14.20 | 12 | -91 | -1 |
| *Right middle occipital gyrus, hMT* |  |  | 13.99 | 27 | -85 | 23 |
| *Right middle occipital gyrus, hMT* |  |  | 13.77 | 33 | -82 | 5 |
| *Hippocampus / thalamus* | 0.0092 | 3 | 6.72 | 21 | -31 | 2 |

**Table S3:** Activation for the blue and yellow colorimetric and apparent blue and yellow stimuli separately, all compared to baseline. All four contrast included activation in V1-V4, V3A and (small) parts of the fusiform gyrus. Regions are labeled based on the probabilistic maps from the Anatomy Toolbox for SPM (Eickhoff et al., 2005).

| Yellow real | Cluster |  | Peak | Location |  |  |
| --- | --- | --- | --- | --- | --- | --- |
| | $P_{(FWE-corr)}$ | Size (voxels) | T | X | Y | Z |
| *Right cuneus, V3d, V2d* | $p < 0.0001$ | 1409 | 26.33 | 18 | -94 | 8 |
| *Right inferior occipital gyrus, V3v* |  |  | 23.68 | 27 | -91 | -4 |
| *left middle occipital gyrus, LO2* |  |  | 19.91 | -27 | -91 | 5 |
| *Left Rolandic operculum* | 0.00096 | 12 | 7.89 | -45 | -25 | 23 |
| *Left Rolandic operculum* | 0.024 | 1 | 6.71 | -42 | -7 | 17 |
| *Right temporal middle gyrus* | 0.024 | 1 | 6.62 | 42 | -64 | 5 |
| *Right fusiform gyrus* | 0.024 | 1 | 6.62 | 30 | -58 | -13 |
| Blue real | Cluster |  | peak | location |  |  |
| | $P_{(FWE-corr)}$ | Size (voxels) | T | X | Y | Z |
| *Right inferior occipital gyrus, LO2* | $p < 0.0001$ | 469 | 20.67 | 27 | -91 | -1 |
| *Right superior occipital gyrus, cuneus* |  |  | 19.21 | 18 | -94 | 8 |
| *Left occipital middle gyrus, LO2* | $p < 0.0001$ | 538 | 17.41 | -24 | -94 | 5 |
| *Left lingual gyrus, fusiform gyrus, V4* |  |  | 16.85 | -21 | -85 | -10 |
| *Left occipital middle gyrus, V3d* |  |  | 16.28 | -18 | -97 | 11 |
| *Left Rolandic operculum* | 0.025 | 1 | 6.36 | -42 | -7 | 17 |
| Apparent Yellow | cluster |  | peak | location |  |  |
| | $P_{(FWE-corr)}$ | Size (voxels) | T | X | y | Z |
| *Right cuneus, superior occipital gyrus, V3d* | $p < 0.0001$ | 1648 | 18.63 | 18 | -94 | 11 |
| *right inferior occipital gyrus, Calcarine sulcus, V3v, V4* |  |  | 18.57 | 27 | -91 | -1 |
| *Left inferior/middle occipital gyrus, Calcarine sulcus, V3v, V4* |  |  | 16.50 | -24 | -88 | -10 |
| Apparent Blue | Cluster |  | Peak | location |  |  |
| | $P_{(FWE-corr)}$ | Size (voxels) | T | X | Y | Z |
| *Right cuneus* | $p < 0.0001$ | 1368 | 21.67 | 18 | -94 | 11 |
| *Left fusiform gyrus, V3d* |  |  | 14.99 | -18 | -94 | 14 |
| *V1* |  |  | 14.41 | 12 | -91 | 2 |

**Table S4:** Differential activation for colorimetric and apparent stimuli. Regions are labeled based on the probabilistic maps from the Anatomy Toolbox for SPM (Eickhoff et al., 2005). Apparent colors showed higher activation in more anterior parts of the early visual cortex and the fusiform gyrus mostly.

| **Colorimetric > apparent colors** | cluster |  | peak | Location |  |  |
| --- | --- | --- | --- | --- | --- | --- |
| | $p_{(FWE-corr)}$ | Size (voxels) | T | X | Y | z |
| *Right middle occipital gyrus, V3d* | $p < 0.0001$ | 11 | 9.70 | 27 | -94 | 11 |
| **apparent > colorimetric colors** | cluster |  | peak | Location |  |  |
| | $p_{(FWE-corr)}$ | Size (voxels) | T | x | y | z |
| *Left lingual gyrus, V1* | $p < 0.0001$ | 183 | 10.41 | -6 | -76 | 5 |
| *Left cuneus, V3d* |  |  | 7.53 | 3 | -76 | 26 |
| *Left calcarine sulcus, V1* |  |  | 7.14 | -6 | -91 | 8 |
| *Right fusiform gyrus* | $p < 0.0001$ | 20 | 8.06 | 30 | -55 | -10 |
| *Left fusiform gyrus* | 0.00033 | 12 | 7.87 | -27 | -61 | -10 |
| *left fusiform gyrus* | 0.00059 | 10 | 7.62 | -30 | -70 | -13 |
| *Right calcarine sulcus, V1* | 0.011 | 2 | 7.12 | 15 | -82 | 2 |
| *Left lingual gyrus* | 0.0045 | 4 | 7.06 | -21 | -49 | -10 |
| *Left cuneus, V3d* | 0.0045 | 4 | 7.04 | -6 | -91 | 29 |
| *Right lingual gyrus* | 0.002 | 6 | 7.03 | 18 | -43 | -7 |
| *Superior parietal lobule* | 0.019 | 1 | 6.68 | 0 | -82 | 35 |
| *Right lingual gyrus, V3v* | 0.011 | 2 | 6.64 | 18 | -70 | -7 |

### Decoding of apparent color (trial-wise regressors)

Apparent yellow and blue could be discriminated in the activity patterns of V1, V2, V3 and V4 (V1: 58.89%, p_corr_ = 0; V2: 59.59%, p_corr_ = 0; V3: 60.23%, p_corr_ = 0; V4: 58.32%, p_corr_ = 0; Figure S2A). These higher decoding accuracies for apparent colors than for colorimetric colors might at first sight seem surprising, however, given that it was not possible to separate out the center and surround fully, these classifications could be based on the color of the illuminant in the surround, which was physically presented. Although this would be the opposite color to the one we are trying to decode, this would be consistently the case in both training and testing, and hence could lead to high discriminability.

### Decoding of apparent lightness

Training on apparent lightness and testing on apparent lightness gave decoding accuracies significantly higher than chance in all four ROIs (V1: 61.00%, p_corr_ = 0; V2: 63.68%, p_corr_ = 0; V3: 59.03%, p_corr_ = 0; V4: 56.00%, p_corr_ = 0; Figure S2B). However, as was the case for apparent color, this might be explained by the effect of the perceptually different surround.

**Figure S2:**
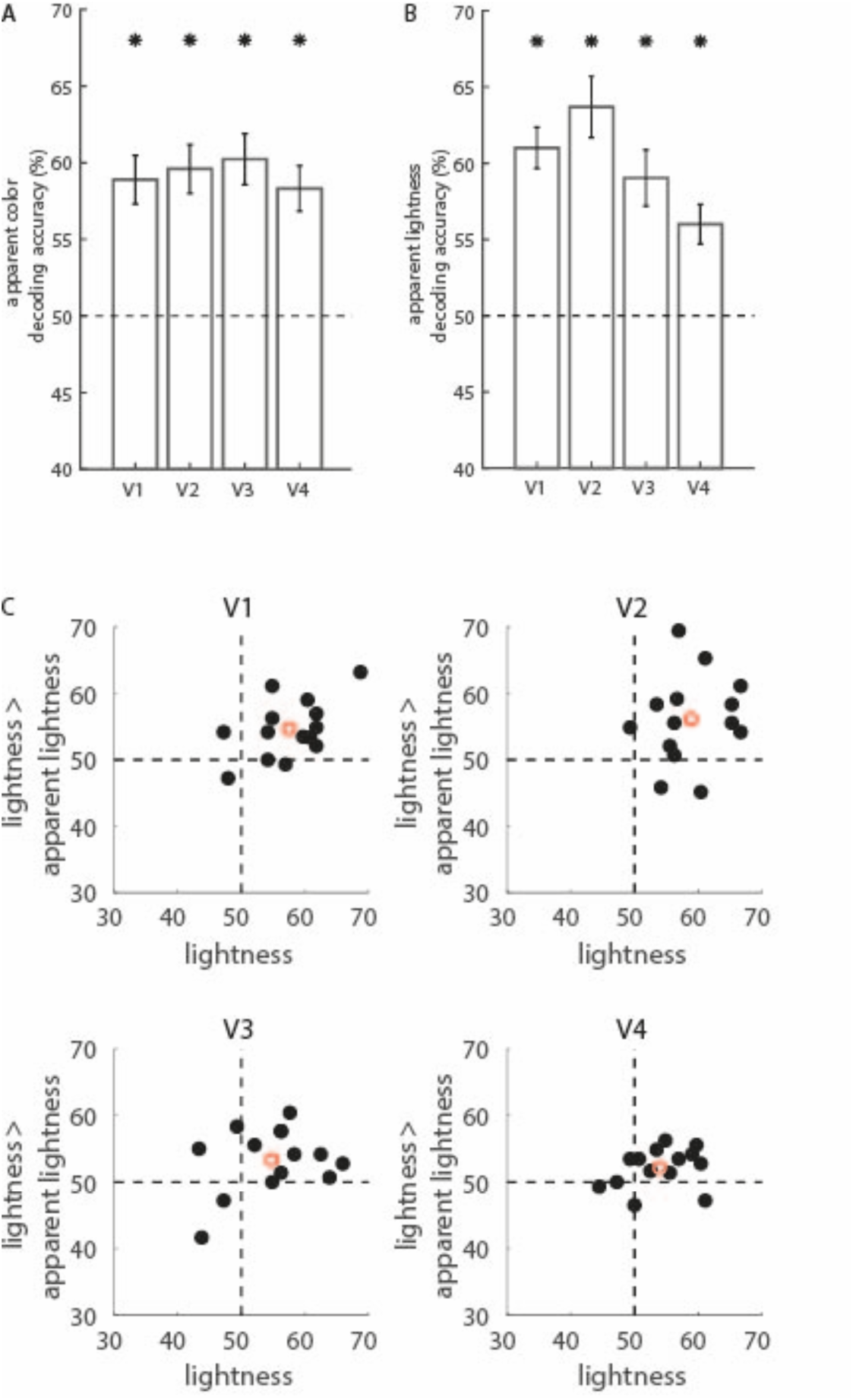
Decoding of apparent color and apparent lightness. A) Decoding accuracy for apparent color. Bars indicate average decoding accuracy for each condition and ROI. Black dashed line indicates the theoretical chance level (50%). Black asterisks indicate p < 0.05 corrected for multiple comparisons, obtained from the permutation tests. B) Decoding accuracy for apparent lightness. Bars indicate average decoding accuracy for each condition and ROI, as in A). C) Lightness decoding versus lightness constancy, for V1-V4. Black dots indicate the data for each individual participant. The red circle is the mean decoding accuracy. Black dashed lines indicate chance level.

### Decoding of colorimetric and color-constant colors in V1-V4 using the run-wise regressors

Colorimetric yellow and blue could be discriminated in V1, V2 and V3, while in V4 there was a trend towards significance (V1: 58.52%, *p_corr_* = 0; V2: 59.05%, *p_corr_* = 0.006; V3: 60.30%, *p_corr_* = 0; V4: 53.45%, *p_corr_* = 0.074; Figure S3B).

To test for color-constant representations in early visual regions, we trained the classifier on the colorimetric colors and tested on the apparent colors, because colorimetric representations of blue and yellow should be present in both cases. Discrimination of colorimetric yellow and blue was numerically possible in V1, V2, V3 and V4, but only reached significance in V2 (V1: 53.18%, *p_corr_* = 0.195; V2: 57.66%, *p_corr_* = 0; V3: 53.0%, *p_corr_* = 0.195; V4: 52.02%, *p_corr_* = 0.198, Figure S3B). Uncorrected, V1 and V3 also showed a trend toward significant generalization. Interestingly, these results are the opposite of what was found using the trial-wise regressors, corroborating the idea that the effect is somewhat present in all four early visual regions, but relatively weakly and hovering around significance.

**Figure S3:**
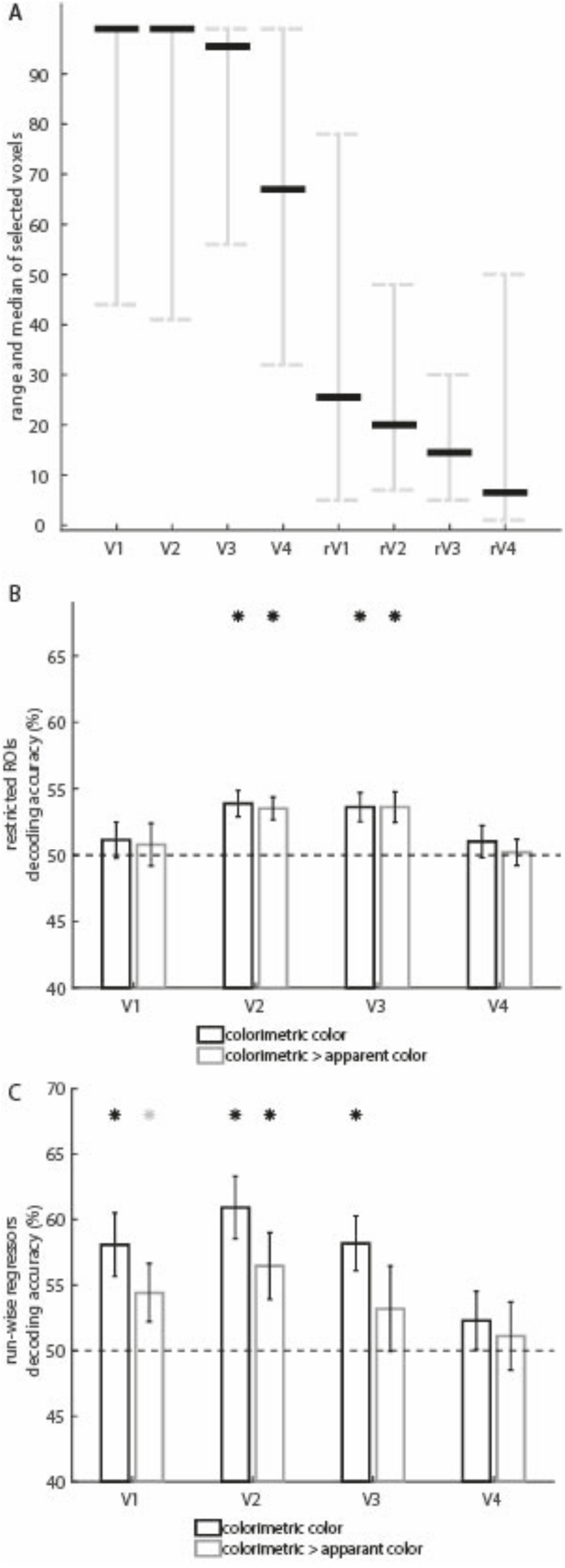
Control analyses. C) Range and median number of voxels selected per ROI based on center activation (V1-V4), and when excluding surround activation (restrictive, rV1-rV4). Black lines indicate the median number of voxels, gray bars indicate the range of voxels selected in the 18 participants). When excluding voxels that respond to the surround localizer, few voxels are left in each ROI. B) Decoding accuracies (±SEM) for colorimetric (black) and color-constant (gray) colors in V1-V4 when using the restrictive ROIs. Color constant colors were obtained by training the classifier on the colorimetric colors, while testing on the apparent colors. Black dashed line indicates the theoretical chance level (50%). Black asterisks indicate p < 0.05 corrected for multiple comparisons, obtained from the permutation tests, gray asterisks indicate *p* < 0.05 uncorrected. C) Decoding accuracies when using run-wise regressors instead of trial-wise regressor; as in B).

### Lightness constancy and color constancy – full results

Full results can be found in Figure S4.Training on apparent lightness and testing on apparent color was possible in all four ROIs; the same was true for training on apparent color induction and testing on apparent lightness. This is in line with the strong decoding results within apparent color (Figure S2A) and most likely has to do with perceptual input from the surround. Training on lightness and testing on colorimetric blue and yellow was possible in V1, V2 and V3. The reverse, training on colorimetric blue and yellow and testing on lightness was possible only in V3, suggesting that lightness information could be used to discriminate colorimetric yellow and blue, whereas the classifier trained on yellow and blue did not simply rely on lightness differences. Finally, training on apparent lightness and testing on colorimetric yellow and blue was not possible in either ROI, while training on apparent color and testing on lightness gave some significant results in V4.

**Figure S4:**
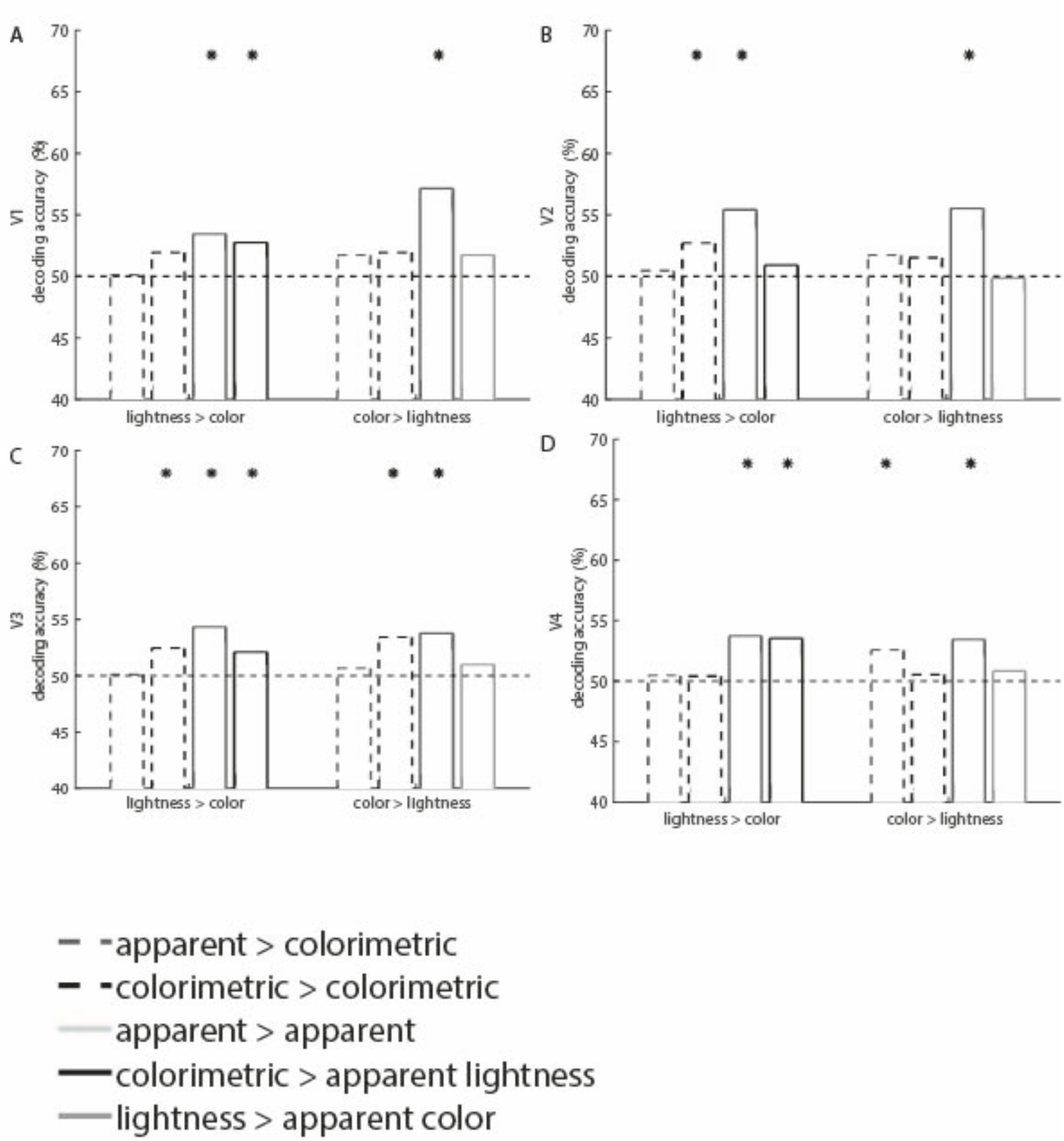
all possible combinations of training colored stimuli (colorimetric and apparent) and testing their gray lightness controls (lightness and apparent lightness), and vice versa. On the left of each plot, training on lightness or apparent lightness controls and testing on colorimetric or apparent color stimuli, on the right of each plot, training on colorimetric or apparent color stimuli and testing on lightness and apparent lightness controls. Dashed line indicates chance level; asterisks indicate significant decoding, uncorrected for multiple comparisons. A) Results for V1; B) results for V2; C) results for V3; D) results for V4.

**Figure S5:**
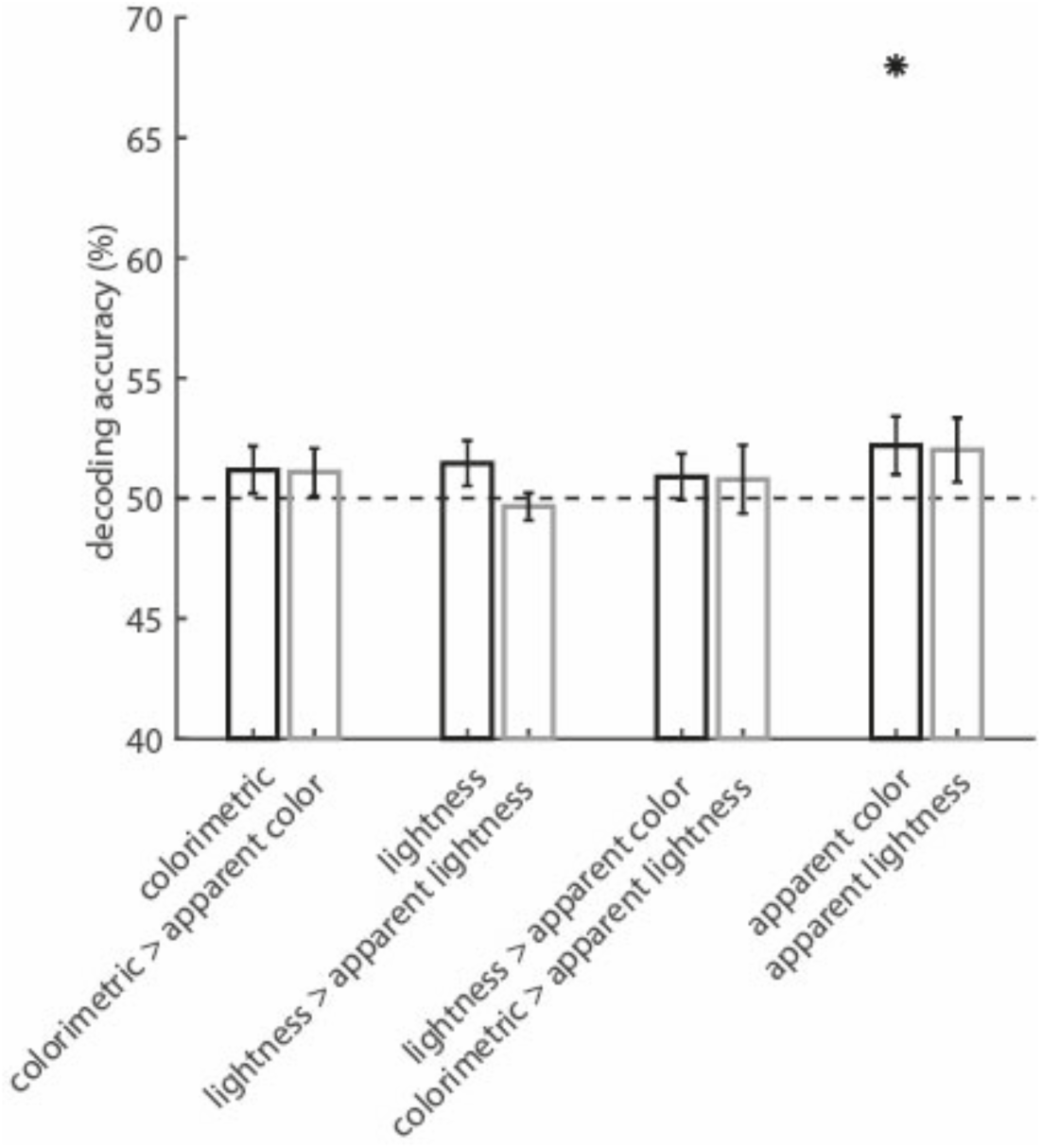
Decoding results in the Rolandic Operculum. Bars indicate average decoding accuracy for each condition; conditions are written below and sorted by analyses. Black dashed line indicates the theoretical chance level (50%). Black asterisks indicate p < 0.05 corrected.

